# Conflict Processing in Schizophrenia: Dissociable Neural Mechanisms Revealed by the N2 and Frontal Midline Theta

**DOI:** 10.1101/2020.07.19.211219

**Authors:** Xiangfei Hong, Fuzhong Yang, Jijun Wang, Chunbo Li, Mingzhou Ding, Jianhua Sheng

## Abstract

Deficits in executive control have long been regarded as one of the hallmark cognitive characteristics in people with schizophrenia (SZ), and current neurocognitive models of SZ generally regard the dysfunctional anterior cingulate cortex (ACC) as the possible neural mechanism. This however, contrasts with recent studies showing that conflict processing, a key component of executive functions that relies on ACC, remains relatively intact in SZ. The current study aimed to investigate this issue through two well-known electrophysiological signatures of conflict processing that have been suggested to origin from ACC, i.e., the N2 component of event-related potentials (ERPs) and frontal midline theta (FMθ) oscillations. We recorded 64-channel scalp electroencephalography from 29 SZ (17 women; mean age: 30.4 years) and 31 healthy control subjects (HC; 17 women; mean age: 29.1 years) performing a modified flanker task. Behavioral data revealed no significant differences in flanker conflict effects (lower accuracy and longer reaction time in incongruent trials than in congruent trials) between HC and SZ. Trial-averaged ERP and spectral analysis suggested that both N2 and FMθ were significantly impaired in SZ relative to HC. Furthermore, by sorting incongruent trials according to their reaction times within individual subjects, we found that the trial-by-trial modulation of N2 (larger amplitude and longer latency in slower trials) which was observed and localized in ACC in HC was totally absent in SZ. By contrast, the trial-by-trial modulation of FMθ (larger power in slower trials) was observed and localized in ACC in both groups, despite a relatively smaller magnitude in SZ. Notably, such trial-by-trial FMθ modulation was only present in incongruent trials that demanded conflict processing. Taken together, our results not only support the idea that FMθ, not N2, serves as the neural substrate of conflict processing in SZ, but also provide novel insights into the functional roles of ACC during executive control, enriching the current neurocognitive models of SZ.

## 1. Introduction

Executive control deficits are one of the hallmark cognitive characteristics in schizophrenia (SZ) (Fioravanti et al., 2005; Barch et al., 2009; Savla et al., 2012). Understanding the neural mechanisms contributing to the deficits is a key step toward developing targeted approaches of treatment and intervention (Carter et al., 2008). Past neuroimaging studies have consistently reported disrupted neural activity in anterior cingulate cortex (ACC), the key brain region involved in executive control (Carter et al., 1998; Botvinick et al., 2004; Gasquoine, 2013), during cognitive tasks requiring executive control (Carter et al., 1997; Carter et al., 2001; Ford et al., 2004; Kerns et al., 2005; Minzenberg et al., 2009). Instead of conceptualizing executive functions coarsely as one unitary cognitive faculty, recent studies that focus on specific components of executive functions have suggested that conflict processing, a core component of executive functions which allows the individual to select a source of information or behavioral tendency in disfavor of distracting or competing sources or tendencies, might remain relatively intact in SZ. Specifically, the behavior of conflict is usually indexed as poorer performance (i.e., slower responses) in incongruent trials (the flankers are associated with opposite responses) than in congruent trials (the flankers are identical to the central target stimuli) in the flanker task (Eriksen and Eriksen, 1974), something referred to as the flanker conflict effect. Increasing studies (Kopp et al., 1994; Kopp and Rist, 1999; Yucel et al., 2002; Lozano et al., 2016; Smid et al., 2016; Ettinger et al., 2018), including a meta-analysis based on data from 1029 SZ and 848 healthy control subjects (HC) (Westerhausen et al., 2013), have reported no significant differences in the flanker conflict effect between SZ and HC. This finding, however, contrasts with the classical neurocognitive models of SZ which generally regard the ACC dysfunction as the neural substrate of executive control deficits (Minzenberg et al., 2009; Lesh et al., 2011).

In parallel with neuroimaging studies, previous electrophysiological investigations have consistently identified two neural signatures of conflict processing, i.e., the N2 event-related potential (ERP) component in the time-domain (Van Veen and Carter, 2002; Folstein and Van Petten, 2008; Ullsperger et al., 2014), and frontal midline theta (FMθ, 4-8 Hz) oscillations in the frequency-domain (Cohen and Donner, 2013; Cohen, 2014b). Since abundant evidences have suggested that both N2 and FMθ have their origin, at least partly, in ACC (Van Veen and Carter, 2002; Wang et al., 2005; Nigbur et al., 2011; Iannaccone et al., 2015; Tollner et al., 2017), one might naturally expect impaired N2 and FMθ in SZ relative to HC, as reported in some recent studies (Neuhaus et al., 2007; Boudewyn and Carter, 2018a; Ryman et al., 2018; Albrecht et al., 2019). For example, one recent study reported impaired trial-averaged FMθ power in SZ than in HC during an AX Continuous Performance Test (CPT) task, suggesting disrupted FMθ during cognitive control in SZ (Ryman et al., 2018). However, it is still difficult to infer the neural mechanisms without assessing the brain-behavior associations, especially at the within-subject level. Indeed, the functional roles of N2 and FMθ in conflict processing in healthy subjects have received indispensably important empirical evidences that both N2 (Yeung et al., 2004) and FMθ (Cohen and Cavanagh, 2011; Cohen and Donner, 2013) were systematically modulated as a function of reaction time (RT) at the within-subject level. In other words, it is still not clear to what extent the impaired N2 and FMθ revealed by trial-averaged analysis in SZ relative to HC reflect the impaired neural responses that are specific to conflict processing or reflect a general impairment of ERPs and neural oscillations in SZ (Uhlhaas and Singer, 2010).

The present study aimed to investigate possible neural substrates of conflict processing in SZ by examining the association between N2/FMθ and flanker conflict effects. We collected 64-channel scalp electroencephalography (EEG) from 30 SZ and 31 HC performing a modified flanker task. In addition to the conventional trial-averaged analysis, we examined trial-by-trial modulations of N2/FMθ by sorting the incongruent trials according to their RTs within each individual. We expected that the trial-by-trial modulations of N2/FMθ might be preserved in SZ, and that if this were the case, it might serve as the neural substrate of conflict processing, and we would further expect to localize their sources in ACC. In particular, the trial-by-trial modulations of N2 and FMθ might be dissociated if they were functionally independent and differentially impacted in SZ. This analysis would provide novel insights for the current neurocognitive models of SZ that have been mainly supported by neuroimaging studies (Minzenberg et al., 2009; Lesh et al., 2011).

## 2. Material and Methods

### 2.1 Participants

The experimental protocol of this study was compliant with the Declaration of Helsinki and approved by the Institutional Review Board of Shanghai Mental Health Center (No. 2017-05R). Informed consent was obtained from each participant before participation. Thirty patients with schizophrenia (SZ) were recruited from the ward of general psychiatry at Shanghai Mental Health Center. The diagnosis was according to the Diagnostic and Statistical Manual of Mental Disorders, Fourth Edition (DSM-IV) and was finally reached at a consensus conference chaired by the director of the ward (J. S.). Thirty-one healthy control subjects (HC) in absence of Axis I disorder according to the Chinese version of the Mini International Neuropsychiatric Interview were recruited locally. Exclusion criteria for both groups included a history of (1) brain injuries, or major medical or neurological disorders, (2) electroconvulsive therapy or magnetic seizure therapy within the recent 6 months, (3) substance abuse disorders within the recent 6 months, or (4) at pregnancy or lactation at the time of the experiment. Additional exclusion criteria for SZ included the comorbidity of other Axis I disorder (not SZ) diagnosed according to the DSM-IV. Additional exclusion criteria for HC included a history or family history (first- or second-degree relative) of psychiatric disorders. All SZ were taking antipsychotics medication and were clinically stable at the time of the experiment. The medication included typical antipsychotics (*N* = 6), atypical antipsychotics (*N* = 29) or both (*N* = 5). Additional medications in SZ included antiparkinson medication (*N* = 4), antidepressants (*N* = 1), anxiolytics (*N* = 1) and antiepileptics (*N* = 1). Positive and Negative Syndrome Scale (PANSS) and Clinical Global Impression were used to assess the symptom severity for SZ. One SZ was excluded in the final analysis due to the difficulty in understanding the experimental task. The demographical information and clinical measurements of the final sample (29 SZ, 31 HC) are summarized in Table 1.

**Table 1.** Group demographics and clinical measures (mean ± SD).

|  | HC (N = 31) | SZ (N = 29) | *p*-value |
| --- | --- | --- | --- |
| *Demographics* |  |  |  |
| Age | 29.1 ± 10.4<br>(range 18-57) | 30.4 ± 8.7<br>(range 19-48) | 0.599<sup>1</sup> |
| Male/Female | 14/17 | 12/17 | 0.768<sup>2</sup> |
| Education (years) | 14.4 ± 2.7 | 12.9 ± 3.1 | 0.054<sup>1</sup> |
| Left-/Right-handed<sup>3</sup> | 1/30 | 0/29 |  |
| *Clinical measures* |  |  |  |
| Age of symptom onset (years) | | 20.2 ± 5.1 | |
| Length of current hospital stay (days) | | 56.4 ± 129.5 | |
| Clinical Global Impression | | 4.7 ± 1.0 | |
| PANSS-positive | | 20.3 ± 8.6 | |
| PANSS-negative | | 18.6 ± 5.6 | |
| PANSS-general | | 38.7 ± 13.0 | |
| PANSS-total | | 77.6 ± 22.4 | |

<sup>1</sup> Independent-samples *t*-test
<sup>2</sup> Chi-square test
<sup>3</sup> Assessed by Edinburgh Handedness Inventory

### 2.2 Stimuli and procedures

Subjects performed a modified flanker task in the experiment (Eriksen and Eriksen, 1974; Bunge et al., 2002; Iannaccone et al., 2015). As shown in Fig. 1, an array of five stimuli was first presented in each trial, which included a center arrow (target, visual angle 0.8°) and two stimuli on either side (flankers, 1°). The entire array of stimuli subtended 6.6° horizontally and 1° vertically, which was consistent with previous study for generating a robust flanker effect (Bunge et al., 2002). There were three types of Go trials (Congruent-Go or CON-Go, Incongruent-Go or INC-Go, Neutral-Go or NEU-Go) and one type of No-Go trials. During Go trials, subjects were asked to press a button using the index finger of the left hand if the center arrow pointed to the left, and press another button using the index figure of the right hand if it pointed to the right. They were required to respond as quickly and as accurately as possible. In CON-Go trials, the flankers were arrows pointing to the same direction as the target. In INC-Go trials, the flankers were arrows pointing to the opposite direction of the target. In NEU-Go trials, the flankers were diamonds and not associated with responses. In No-Go trials, the flankers were X’s, indicating that subjects should withhold their responses. All types of trials were presented in random sequence with equal probability (25% each). In each trial, the stimulus was presented for 800 ms, followed by a random inter-trial interval with blank screen between 800-1200 ms.

**Figure 1.**
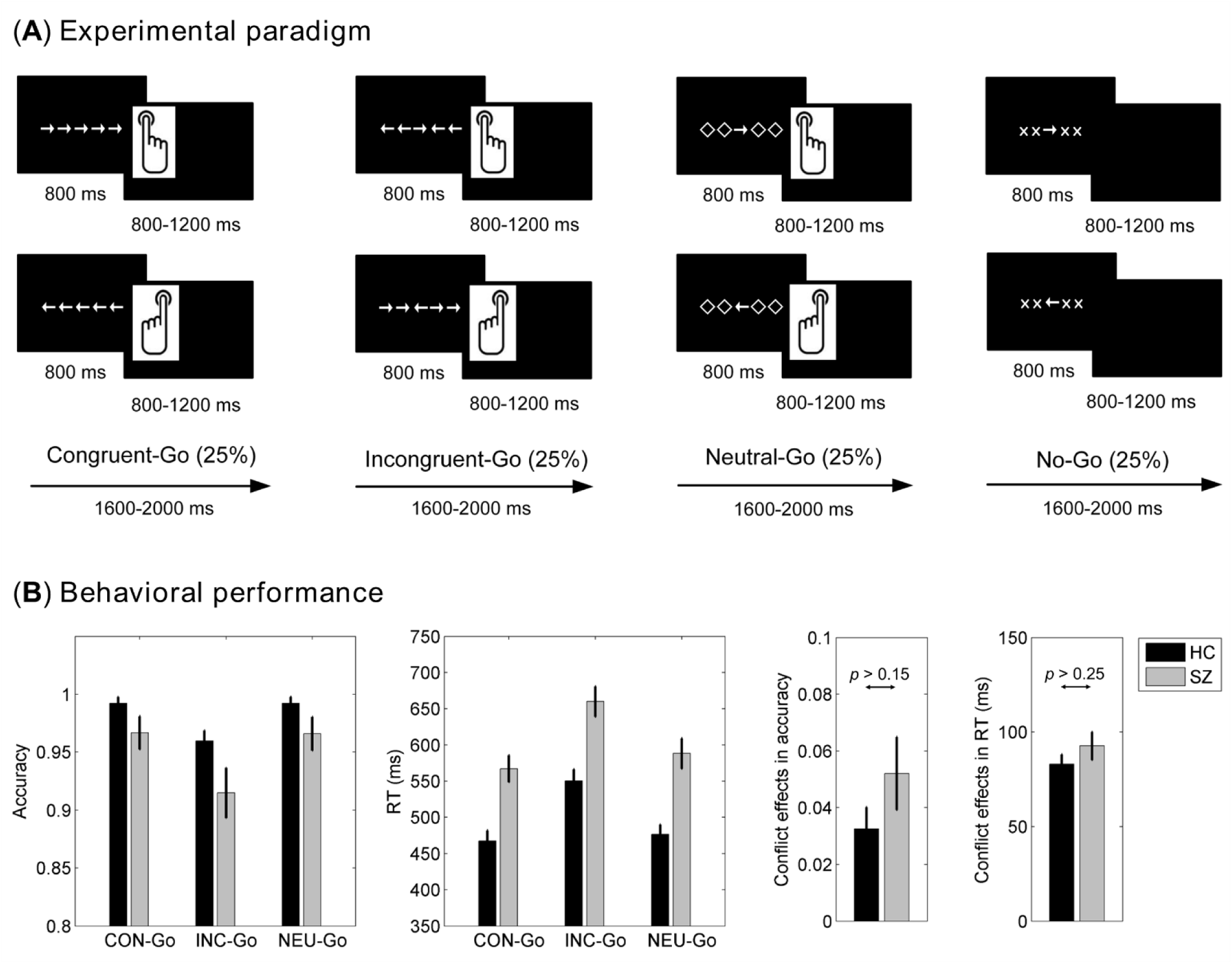
Experimental paradigm and behavioral performance. (*A*) In each trial, subjects viewed an array of stimuli on the screen and responded by pressing the left button when the central arrow pointed to the left and by pressing the right button when it pointed to the right. In Congruent-Go trials (CON-Go), the flankers were arrows pointing to the same direction as the central arrow. In Incongruent-Go trials (INC-Go), they were arrows pointing to the opposite direction as the central arrow. In Neutral-Go trials (NEU-Go), they were diamond-shaped stimuli and not associated with a response. Subjects should refrain from pressing a button in No-Go trials, when the flankers were X’s. (*B*) Accuracy and reaction times (RTs) were shown for CON-Go, INC-Go and NEU-Go trials in each subject group. Flanker conflict effects were quantified as the differences in accuracy (CON-Go *minus* INC-Go) and RT (INC-Go *minus* CON-Go) between CON-Go and INC-Go trials. The flanker conflict effects were not significantly different between the two groups. See main text for the details about statistics. Vertical bars indicate mean ± SEM.

The experimental design was implemented in E-Prime (Version 2.0, Psychology Software Tools, Inc., Sharpsburg, USA), and responses were recorded through the response box included in the E-Prime toolkit. All stimuli were presented on a 19-inch LCD (1280 x 1024 resolution) monitor positioned 60 cm in front of the subject. Each block consisted of 100 trials, lasting for ∼3 min. Subjects were first given the experimental instructions, then trained for at least one block to get familiar with the experiment. After that, each subject finished 6 blocks (600 trials in total, 150 trials per condition), with a short break (1-2 min) between successive blocks.

### 2.3 EEG recording and preprocessing

Continuous EEG signals were recorded during the experiment using the BrainAmp DC amplifier and BrainCap with Fast’n Easy 64Ch standard layout (Brain Products GmbH, Gilching, Germany). EEG signals were recorded through 63 scalp electrodes (Fp1, Fp2, F3, F4, C3, C4, P3, P4, O1, O2, F7, F8, T7, T8, P7, P8, Fz, Cz, Pz, FC1, FC2, CP1, CP2, FC5, FC6, CP5, CP6, FT9, FT10, TP9, TP10, F1, F2, C1, C2, P1, P2, AF3, AF4, FC3, FC4, CP3, CP4, PO3, PO4, F5, F6, C5, C6, P5, P6, AF7, AF8, FT7, FT8, TP7, TP8, PO7, PO8, Fpz, CPz, POz, Oz), with AFz and FCz as recording ground and reference, respectively. The electrode impedance was maintained below 10 kΩ throughout the experiment. EEG signals were digitized at the sampling rate of 1000 Hz with an online anti-aliasing filter (0.016-250 Hz). To monitor eye blinks, one additional electrode (IO) was placed below the right eye, and the difference between electrode IO and Fp2 was calculated as the bipolar VEOG derivation in the offline analysis. The difference between FT9 and FT10 was calculated as the bipolar HEOG derivation.

EEG preprocessing was conducted offline using EEGLAB (Delorme and Makeig, 2004) and ERPLAB toolboxes (Lopez-Calderon and Luck, 2014). Continuous EEG data first went through a two-way band-pass Butterworth filter with zero phase shift (0.1-40 Hz; roll-off slope: 12 dB/oct), followed by a Parks-McClellan notch filter at 50 Hz. After that, EEG data were down-sampled to 250 Hz. Independent component analysis using Infomax algorithm was then performed to correct for ocular artifacts. Corrected EEG data were re-referenced to the algebraic average of the two mastoid electrodes (TP9 and TP10), and the original recording reference electrode was recalculated as FCz in the meantime. Continuous EEG data were segmented into stimulus-related epochs (−1000–1000 ms relative to stimulus onset). It is worth noting that we segmented longer epochs than that we were interested in, so that we can trim the first and last 200 ms data to avoid edge artifacts in time-frequency analysis (see below). Two rounds of artifact detection were performed on each channel for each epoch: (i) the amplitude difference within a moving window (width: 200 ms; step: 50 ms) was examined by the peak-to-peak function (threshold: 100 μV); (ii) the absolute amplitude was examined by a simple voltage threshold function at each time point (threshold: 100 μV). Epochs with artifacts detected on any channel were marked as bad. The trial rejection rate was 8.88% ± 8.07% (mean ± SD) for the HC group, and 14.68% ± 15.41% for the SZ group. Only artifact-free epochs with correct behavioral responses were included in the following analysis. We note that No-Go trials were not analyzed with respect to the main question of interest in the present study.

### 2.4 ERP analysis

Trial-averaged ERP analysis was conducted by averaging EEG epochs of the same trial condition with the 200 ms pre-stimulus interval as baseline, which yielded stimulus-related ERPs for each electrode, trial condition and subject. To isolate conflict-related N2, difference waves (incongruent *minus* congruent) were constructed for each channel and subject. Since conflict-related N2 was typically observed over the middle frontal scalp region, we averaged the ERPs within a middle frontal ROI including Fz, FCz, Cz, FC1 and FC2. The N2 was measured on the difference waves averaged within the middle frontal ROI, and referred to as N2d hereafter. Besides the common mean amplitude approach (measurement window: 380-420 ms for both groups), we also measured the signed area amplitude (negative area) of N2d with a broader measurement window (300-500 ms in both groups), a more robust measurement of amplitudes for difference waves that is less sensitive to the selection of measurement windows (Luck, 2014). The 50% signed area latency approach was applied to measure the latency of N2d (Luck, 2014).

For trial-by-trial ERP analysis, we first sorted all incongruent trials according to their RTs, and equally divided them into three bins of trials with different response speeds, i.e., fast, medium and slow. We then averaged trials in each bin to get ERPs, and constructed difference waves between the ERPs averaged in each trial bin and the ERPs averaged from all congruent trials. This procedure was performed for each subject separately. After that, the signed area amplitude and 50% signed area latency of N2d for each trial bin were measured based on the difference waves averaged within the middle frontal ROI, with 300-500 ms as the measurement window for all three trial bins in both groups.

### 2.5 FMθ analysis

Time-frequency analysis was conducted on EEG epochs using a frequency-domain wavelet convolution approach (Cohen, 2014a). For each EEG epoch, its power spectrum, obtained by the fast Fourier transform, was multiplied by the power spectrum of complex Morlet wavelets, i.e., 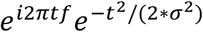, where *t* denotes time, *f* denotes frequency which increases from 2 to 30 Hz in 25 logarithmically spaced steps, and *σ* denotes the width of each frequency band which is 4/2*πf* in this case. The multiplication in the frequency domain is equivalent to the convolution of EEG epoch and Morlet wavelets in the time domain. The complex signal from each convolution was recovered through the inverse fast Fourier transform. The power for specific frequency at each time point was defined as the squared magnitude of the complex signal, yielding the time-frequency power representation for each EEG epoch. For trial-averaged analysis, the power representation was then averaged across all trials of the same condition. This procedure was repeated for each electrode, trial condition, and subject. To make the power comparable between trial conditions and subjects, power values were normalized by the average power during baseline period (−800-−200 ms pre-stimulus) at each frequency using a decibel (dB) tramsform, i.e., dB power = 10 * log10(power/baseline). Theta-band dB power was obtained by averaging normalized dB power within 4-8 Hz and the 300-700 ms post-stimulus interval. Similar to ERP analysis, FMθ power was derived by averaging theta dB power within the middle frontal ROI (Fz, FCz, Cz, FC1 and FC2). For trial-by-trial analysis, we sorted all incongruent trials into three bins using the same approach as that used for ERP analysis, and performed the same time-frequency analysis for each trial bin. To account for the differences in FMθ peak timing between different trial bins (see Fig. 5A), we adjusted the measurement window for FMθ power as 200-600 ms, 250-650 ms and 300-700 ms for fast, medium and slow trials, respectively. For the control purpose, we also performed the same trial-by-trial analysis for congruent trials.

**Figure 2.**
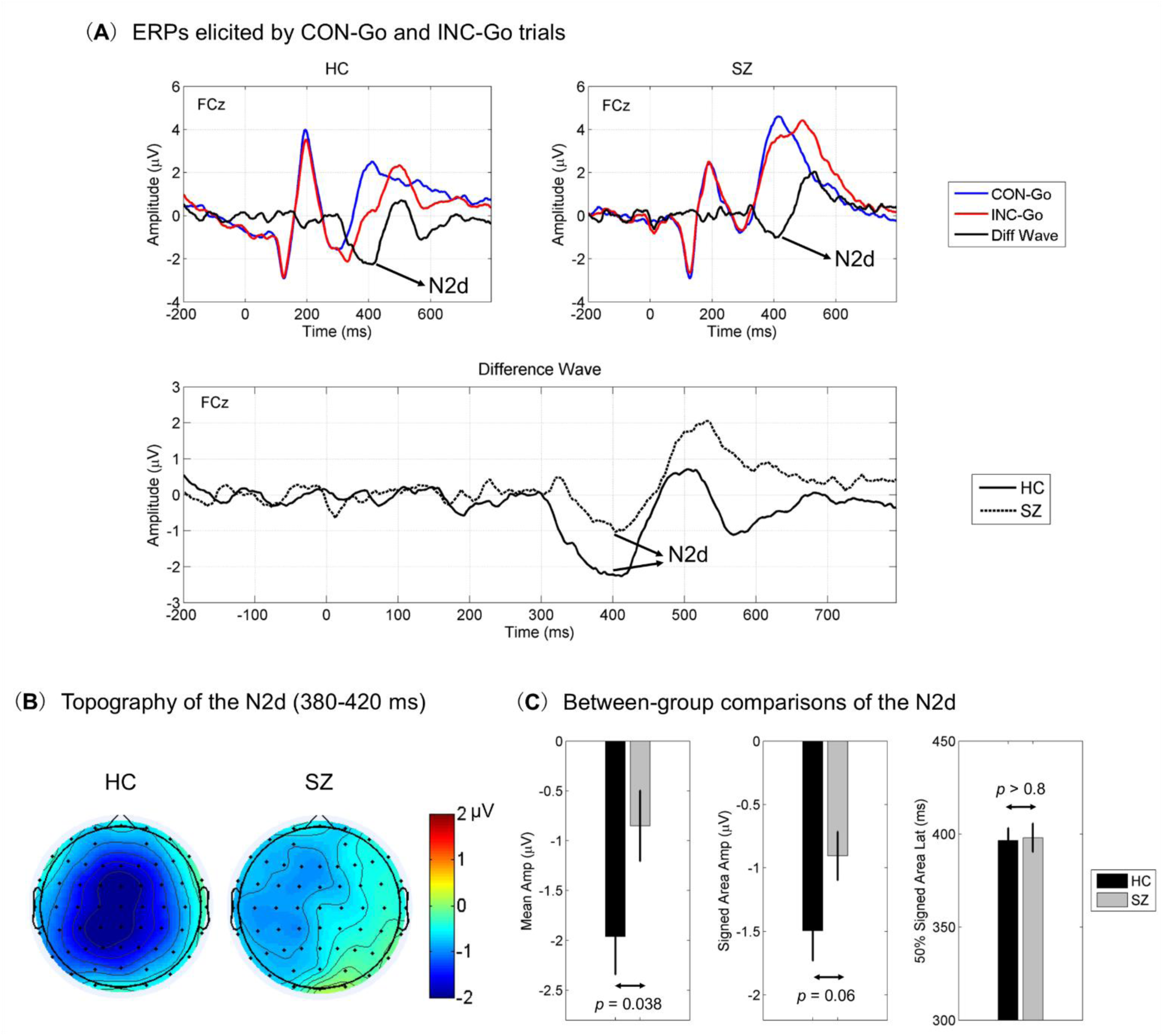
Results of trial-averaged N2d. Grand-averaged difference waves between congruent (CON-Go) and incongruent (INC-Go) trials at electrode FCz (INC-Go *minus* CON-Go) for each group are shown in panel *A*, in which the N2d can be clearly observed. The topographical maps of the N2d averaged within 380-420 ms post-stimulus interval are shown in panel *B*. Panel *C* shows the comparison of N2d amplitude (mean amplitude, signed area amplitude) and latency (50% signed area latency) between the two groups. Vertical bars indicate mean ± SEM.

**Figure 3.**
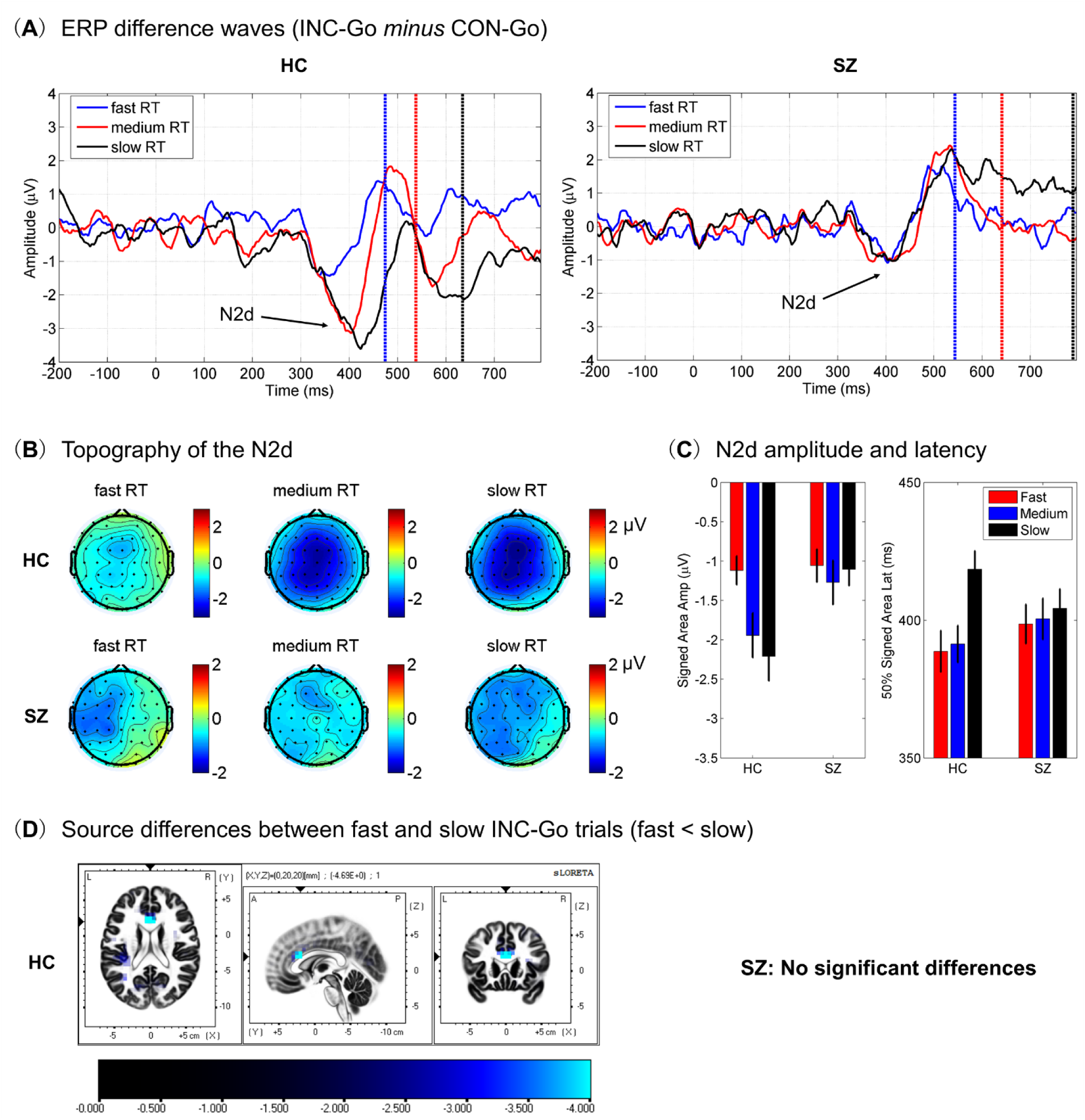
Results of RT-sorted trial-by-trial N2d. Panel *A* illustrates the ERP difference waves between congruent (CON-Go) and incongruent (INC-Go, sorted into fast, medium and slow bins) trials at electrode FCz, with vertical dash lines representing the mean RT in each trial bin. Panel *B* illustrates the topographical maps of N2d, averaged within 325-375 ms for fast trials, 375-425 ms for medium trials and 400-450 ms for slow trials in HC, and within 375-425 ms for all trial bins in SZ. Panel *C* illustrates the signed area amplitude and 50% signed area latency of N2d based on the difference waves averaged within the middle frontal ROI. See main text for the details about statistics for panel *C*. Vertical bars indicate mean ± SEM. Panel *D* shows the source differences between fast and slow trials in each subject group.

**Figure 4.**
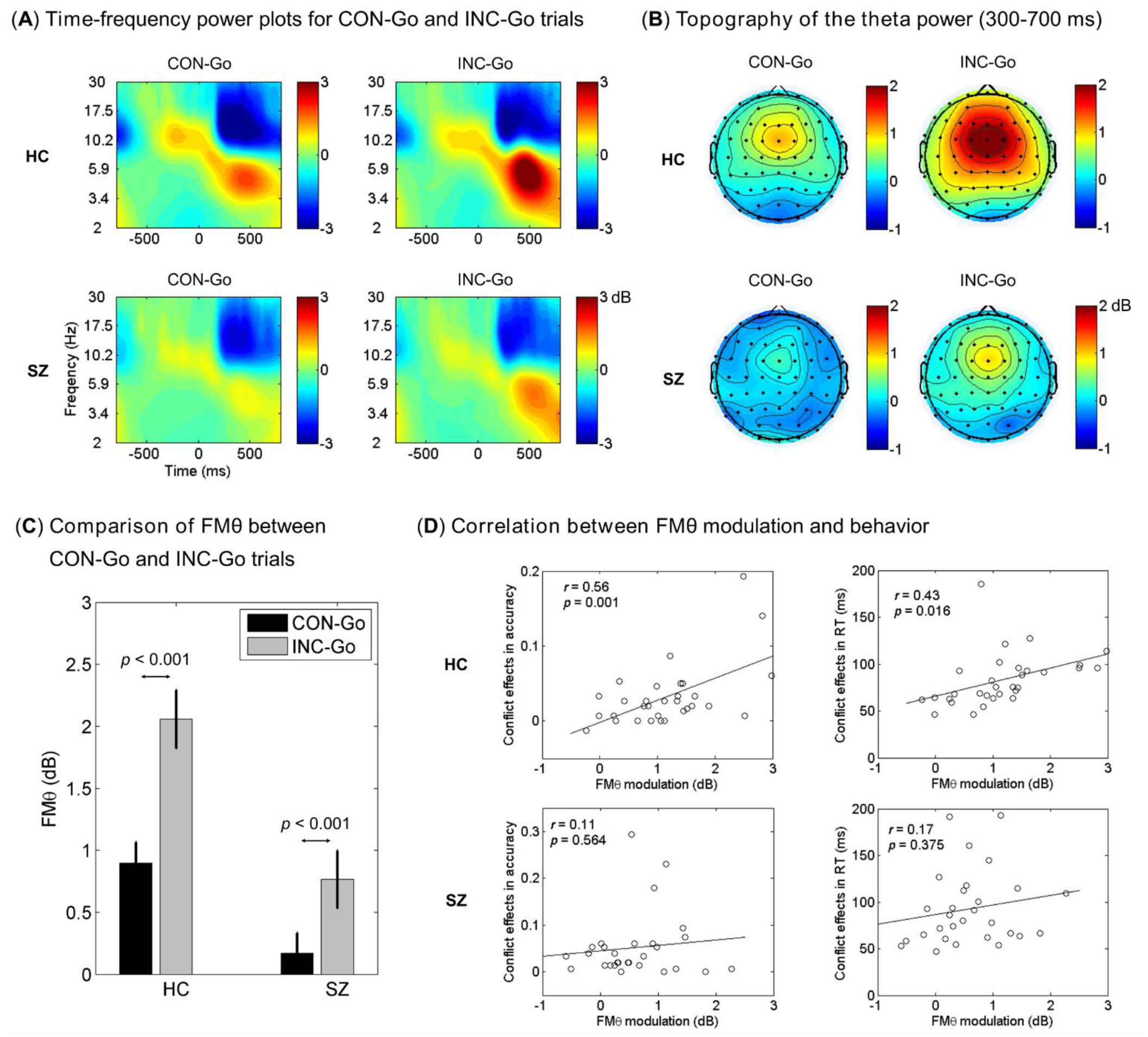
Results of trial-averaged FMθ. Grand-averaged time-frequency power plots at electrode FCz for both congruent (CON-Go) and incongruent (INC-Go) trials are shown in panel *A*, and the topographical maps of theta power averaged within 300-700 ms post-stimulus interval are shown in panel *B*. Panel *C* shows the strength of FMθ power for CON-Go and INC-Go trials. Vertical bars indicate mean ± SEM. Panel *D* shows the correlation between conflict-related FMθ modulation (INC-Go *minus* CON-Go) and behavior in each group. The flanker conflict effects of behavior were defined as the differences in accuracy and RT between CON-Go and INC-Go trials (see Fig. 1B).

**Figure 5.**
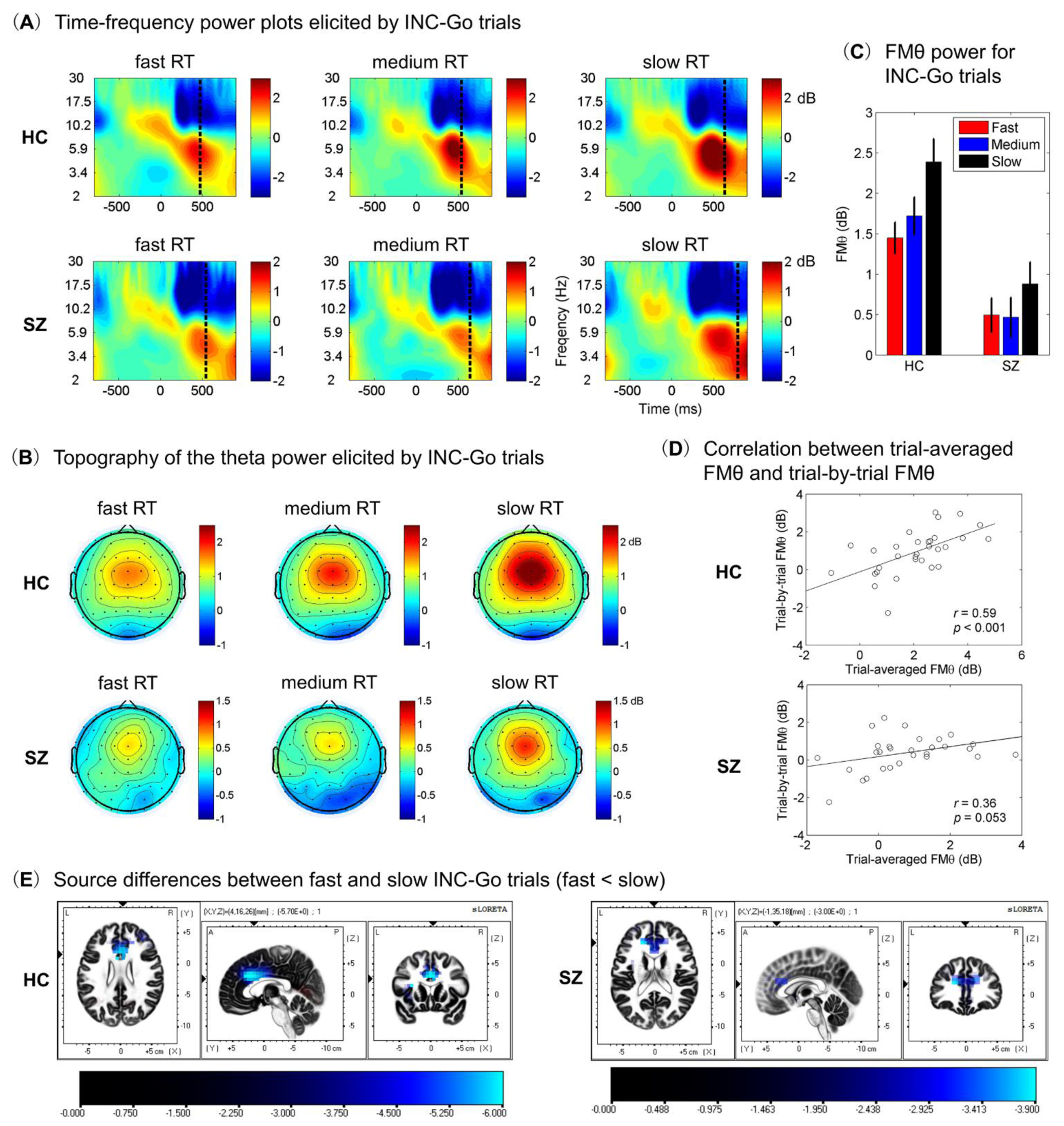
Results of RT-sorted trial-by-trial FMθ. Panel *A* illustrates time-frequency power plots for incongruent (INC-Go, sorted into fast, medium and slow bins) trials at electrode FCz, with vertical dash lines representing the mean RT in each trial bin. Panel *B* illustrates the topographical maps of FMθ power, averaged within 200-600 ms for fast trials, 250-650 ms for medium trials and 300-700 ms for slow trials in both HC and SZ. Panel *C* illustrates FMθ power values averaged within the middle frontal ROI. See main text for the details about statistics for panel *C*. Vertical bars indicate mean ± SEM. Panel *D* shows the correlation between trial-averaged FMθ and trial-by-trial FMθ modulation (slow *minus* fast) in INC-Go trials. Panel *E* shows the source differences of FMθ between fast and slow trials in each subject group.

It is worth noting that FMθ has been suggested to reflect a modulation of ongoing theta oscillations (referred to as “induced” oscillations) that are not phase-locked to the stimulus in conflict processing (Cohen and Donner, 2013; Cavanagh and Frank, 2014; Cohen, 2014b). However, the phase-locked ERP activity, which also includes oscillatory power within the theta-band (referred to as “evoked” oscillations), could contaminate the induced theta-band power obtained by time-frequency analysis (Cohen and Donner, 2013). To minimize the contribution of evoked oscillations, the common approach in the literature was to subtract the averaged ERPs from each EEG epoch before time-frequency analysis. This step was performed for each electrode, trial condition and subject in the present study. However, it has also been suggested that the subtraction of averaged ERPs might further artificially cause residual oscillations due to the trial-by-trial variability of the ERPs (Wang and Ding, 2011). Therefore, we performed an additional set of analyses by subtracting single-trial ERPs estimated by the ASEO (analysis of single-trial ERPs and ongoing-activity) (Xu et al., 2009) from corresponding single-trial EEG epochs before time-frequency analysis. This helps to minimize the possibility that the induced oscillations are driven by variations in single-trial ERPs. In brief, we found that the FMθ power for incongruent trials at electrode FCz was significantly reduced when the averaged ERPs were subtracted than when ERPs were not subtracted in both groups (for HC: 2.40 ± 0.26 dB vs. 2.73 ± 0.27 dB, *t*_(30)_ = −7.056, *p* < 0.001; for SZ: 0.96 ± 0.25 dB vs. 1.20 ± 0.25 dB, *t*_(28)_ = −8.820, *p* < 0.001), suggesting that evoked power was removed by subtracting averaged ERPs, which was consistent with previous study (Cohen and Donner, 2013). However, we observed similar FMθ power between subtracting the averaged ERPs and subtracting the single-trial ERPs in both groups (for HC: 2.40 ± 0.26 dB vs. 2.47 ± 0.27 dB, *t*_(30)_ = −1.025, *p* = 0.313; for SZ: 0.96 ± 0.25 dB vs. 1.03 ± 0.29 dB, *t*_(28)_ = −0.849, *p* = 0.403), suggesting that FMθ oscillations in our data mainly represent the induced oscillations rather than the residual oscillatory activity after subtracting the averaged ERPs. Thus, we only reported the results from subtracting the averaged ERPs in the present study.

### 2.6 Source localization analysis

We applied standardized low-resolution brain electromagnetic tomography (sLORETA) approach to estimate the sources of trial-by-trial modulations of N2d and FMθ. Briefly, sLORETA calculates the standardized current source density for each of the 6239 voxels at 5 mm spatial resolution in the cortical gray matter and the hippocampus normalized to the Montreal Neurological Institute (MNI) space. sLORETA solves the inverse problem by assuming that neighboring neuronal sources should have similar electrical activities (Pascual-Marqui, 2002). In the present study, we computed the source images of N2d averaged within a 40 ms time window centered at the average latency for each trial bin (response speed) and subject: (1) 388 ms (fast), 391 ms (medium) and 418 ms (slow) for the HC group; (2) 398 ms (fast), 400 ms (medium) and 404 ms (slow) for the SZ group. For FMθ, we computed the cross-spectral matrices for each EEG epoch in theta-band (4-8 Hz) during post-stimulus intervals that were adjusted for the timing differences of FMθ (fast: 200-600 ms, medium: 250-650 ms, slow: 300-700 ms). The cross-spectral matrices were then averaged across all epochs in the same trial bin for each subject, and submitted to the source analysis in sLORETA, yielding source images of spectral density at each voxel. Voxel-by-voxel within-subject comparisons of source images were performed to examine possible source differences between different trial bins.

### 2.7 Statistical analysis

Behavioral, scalp ERP and FMθ data were tested in SPSS 22.0 using Repeated-Measures Analysis of Variance (ANOVA), paired-samples *t*-test and independent-samples *t*-test (2-tailed). For ANOVA, Greenhouse–Geisser adjustment was applied to correct the degrees of freedom when the assumption of sphericity was violated. For source localization data, voxel-wise comparisons between different trial bins were performed for the log-transformed source images via paired-samples *t*-tests. Statistical non-Parametric Mapping (SnPM) approach, as implemented in sLORETA software, was used to assess the statistical significance through randomization tests with 5000 permutations. The multiple comparison problem was inherently corrected by the SnPM approach. Pearson correlation (2-tailed) was applied for correlation analysis when necessary. Statistical test was considered as significant when *p* < 0.05. All results were presented as mean ± SEM (standard error of the mean) unless otherwise specified.

## 3. Results

### 3.1 Behavioral results

Accuracy and RT were analyzed using a two-way ANOVA with Trial Condition (congruent vs. incongruent vs. neutral) as a within-subject factor and Subject Group (HC vs. SZ) as a between-subject factor (Fig. 1B). We observed the main effect of Trial Condition for both accuracy (*F*_(2,116)_ = 34.368, *p* < 0.001) and RT (*F*_(2,116)_ = 273.665, *p* < 0.001), indicating a highly significant flanker conflict effect. Moreover, SZ showed an overall decrease of performance, i.e., reduced accuracy and longer RT, compared with HC (Fig. 1B), as indicated by the main effect of Subject Group for both accuracy (*F*_(1,58)_ = 3.756, *p* = 0.058, marginally significant) and RT (*F*_(1,58)_ = 19.716, *p* < 0.001). No significant interaction of Trial Condition × Subject Group was observed for either accuracy or RT, indicating comparable flanker conflict effects between HC and SZ.

We further quantified flanker conflict effects as the differences between congruent and incongruent trials (for accuracy: congruent *minus* incongruent; for RT: incongruent *minus* congruent). As Fig. 1B illustrates, there were no significant differences between the two groups for the flanker conflict effects measured in either accuracy or RT, although SZ showed an increased trend relative to HC. Such results are consistent with the literature, indicating relatively intact conflict processing in SZ compared with HC (Westerhausen et al., 2013).

### 3.2 ERP results

#### 3.2.1 Trial-averaged results

Trial-averaged N2d results are shown in Fig. 2. First, in order to examine whether N2d itself was statistically significant in each group, we compared the mean amplitude of N2d (380-420 ms) against zero.

Note that the signed area amplitude was not used here, because the smallest possible value was zero. Both HC (−1.961 ± 0.381 μV; *t*_(30)_ = −5.148, *p* < 0.001) and SZ (−0.851 ± 0.355 μV; *t*_(28)_ = −2.400, *p* = 0.023) showed significant N2d component. Second, the mean amplitude of N2d was significantly smaller in SZ than that in HC (*t*_(58)_ = −2.125, *p* = 0.038), while this difference became marginally significant when the N2d amplitude was quantified using the signed area approach (HC: −1.496 ± 0.237 μV vs. SZ: −0.906 ± 0.191 μV; *t*_(58)_ = −1.922, *p* = 0.060). Third, no significant difference was observed for the 50% signed area latency of N2d between the two groups (HC: 396.5 ± 6.7 ms vs. SZ: 398.1 ± 7.7 ms; *t*_(58)_ = −0.153, *p* > 0.8). Together, these results indicate that the N2d was not delayed, but was significantly impaired in SZ relative to HC. Finally, we examined brain-behavior relationships by correlating N2d amplitudes (either mean amplitude or signed area amplitude) with flanker conflict effects of behavior (measured in either accuracy or RT). No significant correlations were observed in either group (all *p* > 0.1).

#### 3.2.2 RT-sorted results

RT-sorted trial-by-trial N2d results are shown in Fig. 3. First, the signed area amplitude of N2d was tested by a two-way ANOVA with Response Speed (fast vs. medium vs. slow) as a within-subject factor and Subject Group (HC vs. SZ) as a between-subject factor (Fig. 3C). We observed the main effects of both Response Speed (*F*_(2,116)_ = 6.536, *p* = 0.002) and Subject Group (*F*_(1,58)_ = 4.193, *p* = 0.045), as well as a significant interaction of Response Speed × Subject Group (*F*_(2,116)_ = 4.533, *p* = 0.013). To understand this interaction, we further performed one-way ANOVA with Response Speed (fast vs. medium vs. slow) as the factor in each group. The main effect of Response Speed was observed in HC (*F*_(2,60)_ = 10.702, *p* < 0.001), but not in SZ (*F*_(2,56)_ = 0.410, *p* = 0.666). These results suggest that HC showed increased N2d amplitude as the level of conflict increased. However, such trial-by-trial modulation of N2d was absent in SZ.

Second, we quantified the 50% signed area latency of N2d in different trial bins, which was then tested through the same two-way ANOVA as that used for N2d amplitude (Fig. 3C). We observed the main effect of Response Speed (*F*_(2,116)_ = 5.367, *p* = 0.006) and a marginally significant interaction of Response Speed × Subject Group (*F*_(2,116)_ = 2.701, *p* = 0.071). We thus performed one-way ANOVA with Response Speed (fast vs. medium vs. slow) as the factor in each group. The main effect of Response Speed was observed in HC (*F*_(2,60)_ = 9.253, *p* < 0.001), but not in SZ (*F*_(2,56)_ = 0.206, *p* = 0.815). These results further indicate that the trial-by-trial modulation of N2d was absent in SZ.

Third, the source differences of N2d between fast and slow trial bins were localized in ACC in HC (corrected *p* < 0.05, 1-tailed, fast < slow, see Fig. 3D). By contrast, no significant differences were observed in SZ.

### 3.3 FMθ results

#### 3.3.1 Trial-averaged results

Trial-averaged FMθ results are shown in Fig. 4. FMθ power was submitted to a two-way ANOVA with Trial Condition (congruent vs. incongruent) as a within-subject factor and Subject Group (HC vs. SZ) as a between-subject factor. We observed the main effect of Trial Condition (*F*_(1,58)_ = 83.788, *p* < 0.001), suggesting a significant increase of FMθ in incongruent trials than in congruent trials. The overall strength of FMθ was significantly reduced in SZ than in HC, as indicated by the main effect of Subject Group (*F*_(1,58)_ = 13.403, *p* = 0.001). Moreover, given the significant interaction of Trial Condition × Subject Group (*F*_(1,58)_ = 8.676, *p* = 0.005), we then performed paired-samples *t*-tests between congruent and incongruent trials in each group. Results indicate that FMθ modulation was significant in both groups (both *p* < 0.001, see Fig. 4C). To better understand this interaction, we further quantified the conflict-related FMθ modulation by calculating the difference in FMθ between congruent and incongruent trials (incongruent *minus* congruent). Results suggest that FMθ modulation was significantly greater in HC than in SZ (HC: 1.16 ± 0.14 dB vs. SZ: 0.60 ± 0.12 dB; *t*_(58)_ = 2.946, *p* = 0.005). Finally, we examined brain-behavior relationships by correlating FMθ modulation with flanker conflict effects of behavior (measured in either accuracy or RT). As Fig. 4D illustrates, we observed significant positive correlations in HC, suggesting that subjects with larger flanker conflict effects also showed greater FMθ modulation. Such correlations, however, were absent in SZ.

#### 3.3.2 RT-sorted results

RT-sorted trial-by-trial FMθ results are shown in Fig. 5. FMθ power was tested by a three-way ANOVA with Trial Condition (congruent vs. incongruent), Response Speed (fast vs. medium vs. slow) as within-subject factors, and Subject Group as a between-subject factor. We observed the main effects of Trial Condition (*F*_(1,58)_ = 80.172, *p* < 0.001), Response Speed (*F*_(2,116)_ = 13.910, *p* < 0.001) and Subject Group (*F*_(1,58)_ = 13.004, *p* < 0.001), as well as significant interactions of Trial Condition × Subject Group (*F*_(1,58)_ = 9.939, *p* = 0.003) and Trial Condition × Response Speed (*F*_(2,116)_ = 3.448, *p* = 0.035). These interactions related to Trial Condition suggest different trial-by-trial modulations of FMθ between congruent and incongruent trials. To better understand these interactions, we further performed two-way ANOVA with Response Speed (fast vs. medium vs. slow) as a within-subject factor and Subject Group as a between-subject factor for congruent trials and incongruent trials separately. For incongruent trials, as shown in Fig. 5C, we observed the main effects of both Response Speed (*F*_(2,116)_ = 16.426, *p* < 0.001) and Subject Group (*F*_(1,58)_ = 15.294, *p* < 0.001), as well as a marginally significant interaction of Response Speed × Subject Group (*F*_(2,116)_ = 2.533, *p* = 0.084). Given this interaction, we further performed one-way ANOVA with Response Speed (fast vs. medium vs. slow) as the factor in each group. The main effect of Response Speed was observed in both HC (*F*_(2,60)_ = 15.347, *p* < 0.001) and SZ (*F*_(2,56)_ = 3.562, *p* = 0.035), suggesting that trial-by-trial FMθ modulation was present in both groups. Moreover, the magnitude of trial-by-trial FMθ modulation (slow *minus* fast) was larger in HC than in SZ (HC: 0.94 ± 0.21 dB vs. SZ: 0.39 ± 0.17 dB; *t*_(58)_ = 2.037, *p* = 0.046), which explained the above interaction of Response Speed × Subject Group. By contrast, for congruent trials, we only observed the main effect of Subject Group (*F*_(1,58)_ = 7.761, *p* = 0.007), without other main effects or significant interactions. Such results suggest that trial-by-trial FMθ modulation was only present in incongruent trials that demanded conflict processing, but not in congruent trials in which such demand was minimal.

Furthermore, we examined whether the trial-by-trial FMθ modulation (slow *minus* fast) was associated with the trial-averaged FMθ in incongruent trials. As shown in Fig. 5D, we observed significant positive correlation in HC, i.e., individuals with higher trial-averaged FMθ also showed larger trial-by-trial FMθ modulation. While in SZ, although the correlation strength was reduced relative to HC, the correlation was still marginally significant.

Finally, the source differences of FMθ between fast and slow trial bins were localized in ACC in both HC and SZ (corrected *p* < 0.05, 1-tailed, fast < slow, see Fig. 5E).

#### 3.3.3 Correlating trial-by-trial FMθ modulation with PANSS in SZ

We performed an exploratory analysis for the correlation between trial-by-trial FMθ modulation (slow *minus* fast) and PANSS scores (positive, negative, general, and total scores separately). However, none of these correlations were significant (all *p* > 0.2).

### 3.4 Correlation between N2d and FMθ

To examine whether N2d and FMθ are related, we performed the correlation analysis between trial-averaged N2d amplitude and trial-averaged FMθ power, and between trial-by-trial modulation of N2d amplitude and trial-by-trial modulation of FMθ power. However, none of these correlations were significant in either group (all *p* > 0.2).

## 4. Discussion

The present study investigated possible neural mechanisms of conflict processing in SZ by examining two well-known electrophysiological signatures, i.e., the N2 component and FMθ oscillations. Behavioral and EEG data were recorded from HC and SZ performing a modified flanker task. Behaviorally, we observed comparable flanker conflict effects between HC and SZ, although SZ performed generally worse than HC. EEG results based on trial-averaged analysis suggest that both HC and SZ showed significant conflict effects of N2 and FMθ, and both effects were significantly impaired in SZ. Results based on RT-sorted trial-by-trial analysis at the within-subject level reveal dissociable changes between N2 and FMθ. Specifically, the N2 was significantly greater and delayed when the conflict became larger (indexed by longer RTs) in HC, while this modulation was absent in SZ. The FMθ was also significantly greater when the conflict became larger, and notably, this conflict-related modulation was present in both HC and SZ, but only present in incongruent trials that required conflict processing. Moreover, the magnitude of trial-by-trial FMθ modulation was positively correlated with the magnitude of trial-averaged FMθ, although this correlation was weaker in SZ than in HC. This dissociation indicates that FMθ, not N2, might serve as the neural substrate of intact conflict processing in SZ. Additionally, source localization analysis suggested ACC as the main brain region for trial-by-trial modulations of FMθ, indicating that the function of ACC in SZ was not totally disrupted in executive control.

Although few studies on SZ directly examined the N2 during conflict processing [i.e., flanker task, for research using the attention network test, see Neuhaus et al. (2007), which however, failed to show robust flanker effects of N2 in HC], previous work has shown N2 changes in SZ during response inhibition (Kiehl et al., 2000; Weisbrod et al., 2000; Araki et al., 2016; Krakowski et al., 2016; Hoptman et al., 2018). Notably, the N2 observed in response inhibition tasks has been proposed to reflect the conflict-related processing (i.e., conflict between executing and inhibiting a response) rather than inhibition itself (Nieuwenhuis et al., 2003; Donkers and van Boxtel, 2004), and when combined with our trial-averaged N2 results, it might be inferred that conflict-related N2 is impaired in SZ. Similarly, FMθ, another recently proposed signature for executive control (Cavanagh and Frank, 2014), has also been shown to be substantially impaired in SZ relative to HC during proactive control (Ryman et al., 2018), error processing (Boudewyn and Carter, 2018a), response inhibition (Cooper and Hughes, 2018) and working memory (Schmiedt et al., 2005). In the present study, although SZ showed enhanced FMθ in incongruent trials than in congruent trials, the trial-averaged FMθ modulation was prominently impaired in SZ, which is consistent with previous studies, suggesting that impaired FMθ represents a key deficit of SZ during cognitive control (Boudewyn and Carter, 2018b).

Despite these trial-averaged results that are consistent with the literature, the comparable flanker conflict effects of behavior between HC and SZ still indicate preserved neural substrates of conflict processing in SZ, and our results based on within-subject trial-by-trial analysis suggest FMθ as a candidate for such neural substrates, with evidences from the following three aspects. First, both HC and SZ showed increased FMθ power when the conflict became larger (indexed by longer RTs), albeit to a smaller magnitude of such increase in SZ relative to HC. Second, our results show that FMθ reached its peak before the responses were made, which became more evident when the responses were slower (Fig. 5A). This finding is consistent with previous studies on healthy controls reporting that trial-by-trial fluctuations of FMθ predict RTs in conflict trials (Cohen and Cavanagh, 2011; Cohen and Donner, 2013), suggesting that conflict should be resolved before correct responses. Notably, this predictive relationship between FMθ and RT was preserved in SZ. Third, the positive correlation between trial-averaged FMθ and trial-by-trial modulation of FMθ was also preserved in SZ, despite a weaker strength relative to HC (Fig. 5D).

Although the precise functional role that FMθ might have in the neural implementations of conflict processing is still unclear, modern theories on the computational role of neural oscillations generally suggest that oscillations form a cyclic temporal reference frame for organizing information processing, and more specifically, FMθ may serve to provide such a reference frame by enforcing narrow windows for the detection and signaling of conflict in the neural microcircuit (Wang, 2010; Cohen, 2014b). Consistently, by recoding the single neuron responses and local field potentials in human neurosurgical patients, recent study has shown that processing response conflict might require cross-areal rhythmic neuronal coordination between ACC and dorsolateral prefrontal cortex that is possibly mediated by theta oscillations (Smith et al., 2019). Thus, our results might imply that the FMθ-mediated neuronal computation and communication for conflict processing was functionally preserved in SZ, at least in part, and this might underlie the intact performance of conflict processing.

By contrast, the trial-by-trial modulation of N2 amplitude and latency as a function of conflict (RT), an observation in HC that was consistent with previous research (Yeung et al., 2004), was totally absent in SZ. When combined with FMθ, this pattern of results not only suggests independent roles between N2 and FMθ in conflict processing, but also indicates dissociable changes of these two signatures in SZ, which might be further supported by the absence of correlation between them in either HC or SZ. From the computational perspective, it is worth noting that in the present study, non-phase-locked FMθ oscillations were extracted by time-frequency analysis after subtracting phase-locked ERPs, making them possibly represent independent neural activities (Cohen and Donner, 2013). Notably, we further minimized the possibility that induced FMθ oscillations were driven by variations in single-trial ERPs in this subtraction approach by exacting and subtracting ERPs at the single-trial level.

The source localization analysis in the present study suggests ACC as the main region for trial-by-trial conflict modulations of N2 and FMθ, which is consistent with previous research. However, our findings that the absent N2 modulation while the preserved FMθ modulation naturally warrant a careful revisit for the role of ACC in conflict processing and executive control in SZ. The current neurocognitive models of SZ generally regard the ACC dysfunction as a key contributor for executive control deficits, with the evidences mostly provided by neuroimaging studies (Minzenberg et al., 2009; Lesh et al., 2011). Unfortunately, there is still lack of neuroimaging studies examining the brain-behavior relationship in conflict processing in SZ, especially on the within-subject trial-by-trial basis. The reason might be the difficulty in collecting as many trials as that collected in the electrophysiological studies due to the much slower hemodynamic responses and experimental designs that neuroimaging studies typically rely on. However, by combining source localization and trial-by-trial analysis with a typical EEG task design, our results clearly suggest that ACC is still functionally reactive to the increased demand of conflict processing in SZ, and it is doing so by generating larger FMθ oscillations that might play a key role in the neuronal computation and communication for conflict processing (Cohen, 2014b).

Regarding the experimental paradigm, it has been proposed that the flanker task with constant stimulus– response mappings mostly engage reactive control, which differs from proactive control recruited by AX-CPT and other cue–probe tasks, suggesting impaired proactive control while intact reactive control in SZ (Smid et al., 2016; Ryman et al., 2018). On the other hand, it is worth noting that the preserved performance of conflict processing in SZ might partially reflect the nature of cognitive processes involved in the flanker task. Specifically, it has been proposed that the impairment in executive control is dependent on whether the *inhibition* component is required in the cognitive task (Westerhausen et al., 2013). Evidences supporting this notion come from previous studies that reported significant behavioral impairments in SZ than in HC during the Stroop task (Westerhausen et al., 2011), the Stop-Signal task (Lipszyc and Schachar, 2010), as well as the Go/NoGo task (Kiehl et al., 2000; Weisbrod et al., 2000), all requiring the inhibition of a prepotent, habitual internal or overt response. Consistent with this notion, we also found that SZ showed significantly higher error rates in No-Go trials than HC (results not included here due to the focus of this paper). By contrast, the flanker task, although also requiring the inhibition of incorrect response tendency elicited by incongruent flankers, relies more on the ability to filter out distracting irrelevant information (i.e., flankers around the central target) compared with those inhibition tasks. Notably, it has been suggested that attention filtering in visual modality is likely to be intact in SZ (Luck et al., 2019), and this might aid, at least to some extent, the performance in the flanker task for SZ.

In conclusion, the current study, for the first time, examined both N2 and FMθ during conflict processing and their changes in SZ at both between-subject and within-subject levels. Although conventional trial-averaged analysis suggest that both N2 and FMθ were prominently reduced in SZ, our results based on within-subject trial-by-trial analysis indicate dissociable changes between N2 and FMθ in SZ. The conflict-related trial-by-trial modulations were absent for N2, but preserved for FMθ in SZ, suggesting that FMθ might serve as the possible neural substrate of intact conflict processing in SZ. The source localization analysis of FMθ provides novel evidences that ACC is still functionally reactive under high conflict situations in SZ, enriching the current neurocognitive models of SZ.

## Funding

This work was supported by Research Foundation of Shanghai Municipal Health Bureau (No. 20174Y0020), Shanghai Municipal Natural Science Foundation (No. 18ZR1432700), Research Foundation of Shanghai Jiao Tong University (No. YG2017MS44), National Natural Science Foundation of China (No. 61601294), Shanghai Mental Health Center (2016-QH-01, 2016-YJ-11) and Science and Technology Commission of Shanghai Municipality (No. 13dz2260500). The funding sources had no role in study design, data collection and analysis, decision to publish, or preparation of the manuscript.

## Declarations of Interest

None.

## Notes

### Competing Interest Statement

The authors have declared no competing interest.

